# Striatal cholinergic interneuron pathology and muscarinic signaling independently shape motor dysfunction in a DYT-TOR1A dystonia model

**DOI:** 10.64898/2026.09.13.751056

**Authors:** Caitlin E. Leedy, Jay Li, Samuel S. Pappas, William T. Dauer

## Abstract

Antimuscarinic drugs are among the most effective pharmacological treatments for dystonia, yet the neural substrates underlying their therapeutic effects remain poorly understood. Striatal cholinergic interneurons (ChI) are strongly implicated in dystonia pathophysiology. In a symptomatic mouse model of DYT-TOR1A dystonia, torsinA loss from all striatal neurons causes selective ChI degeneration during juvenile maturation; surviving ChI exhibit persistent morphological, physiological, and connectivity abnormalities. Whether motor dysfunction arises from ChI degeneration, dysfunction of surviving ChI, or altered cholinergic signaling elsewhere in the striatal circuit remains unknown. Here, we distinguish these mechanisms using complementary genetic, pharmacological, and cell-ablation approaches. Surviving ChI became hyperactive coincident with the emergence of neurodegeneration and abnormal movements. Selective prenatal restoration of torsinA in ChI prevented their degeneration and reduced abnormal movements, identifying ChI as an important cellular locus of torsinA-dependent motor dysfunction. Systemic antimuscarinic treatment during juvenile striatal maturation produced persistent behavioral improvement outlasting treatment without preventing ChI degeneration. Direct intrastriatal antimuscarinic administration nearly abolished abnormal movements, identifying the striatum as a critical site of therapeutic action. Surprisingly, extensive ablation of the remaining dorsal striatal ChI neither prevented nor ameliorated abnormal movements and did not diminish the efficacy of systemic or intrastriatal antimuscarinic treatment. Thus, torsinA-dependent ChI pathology contributes causally to motor dysfunction, yet the continued presence of dorsal striatal ChI is not required for either expression of the motor phenotype or its suppression by muscarinic antagonists. These findings dissociate a developmental contribution of ChI pathology from the striatal mechanisms through which antimuscarinic drugs suppress dystonia-related movements.

## Introduction

Dystonia is a movement disorder characterized by abnormal twisting movements, often exacerbated by voluntary or intentional movements [1, 2]. DYT-TOR1A is an early-onset dystonia [3] caused by a dominantly-inherited mutation in the TOR1A gene [4] that impairs the function of torsinA [5–9]. While the precise circuit mechanisms causing abnormal motor behaviors in DYT-TOR1A dystonia have not been fully elucidated, abnormalities within the cholinergic system have been observed in humans, and anticholinergic medications remain a mainstay of treatment for dystonia [10–13]. Diverse cholinergic abnormalities have been observed in animal models of DYT-TOR1A dystonia [14–20], and modulation of cholinergic activity has been demonstrated to induce abnormal movements [21, 22].

DYT-TOR1A is an incompletely penetrant neurodevelopmental disorder and typically manifests before 26 years of age [3]. Evidence from animal models of DYT-TOR1A indicates that torsinA is uniquely important during a critical period of CNS maturation, as torsinA deletion from the CNS of juvenile mice causes neurodegeneration and abnormal limb clasping behaviors but causes no abnormalities when deleted from adult mice [23, 24]. This developmental window coincides with the maturation of striatal circuitry [25–30]. As some of the earliest neurons to migrate to the striatum, cholinergic interneurons (ChI) are uniquely poised to regulate striatal development [29, 31–33].

Striatal ChI contribute to the gating of corticostriatal plasticity by coordinating acetylcholine signaling with cortical activity and phasic dopamine release, thereby determining whether active corticostriatal synapses are strengthened, weakened, or left unchanged [34–39]. Striatal ChI are autonomous pacemakers which provide steady cholinergic tone to the striatum, but exhibit a pause behavior which is hypothesized to signal salient events and contribute to plasticity (reviewed in [40]). This pause develops during the first month of postnatal maturation in mice [26, 41]. Disruptions in striatal cholinergic signaling are implicated in a variety of movement disorders, including dystonia [16, 42, 43]. Abnormal cholinergic tone may disrupt the gating of corticostriatal plasticity, leading to maladaptive basal ganglia output of both the striatonigral and striatopallidal populations of spiny projection neurons (SPNs) [16, 42]. The balance within the striatum between cholinergic and dopaminergic signaling is important for motor learning and proper execution of motor commands [44, 45]. Due to the centrality of striatal ChI to behavior, motor learning, and striatal plasticity, as well as their coincident degeneration during the emergence of clasping in a mouse model of DYT-TOR1A dystonia, we sought to further examine the contribution of striatal cholinergic interneurons to abnormal movement generation.

The “Dlx-CKO” mouse model of DYT-TOR1A dystonia lacks torsinA from forebrain inhibitory and cholinergic interneurons [14]. Similar to the human disorder, Dlx-CKO mice are born phenotypically normal, but develop abnormal limb clasping movements during postnatal maturation that are improved by systemic antimuscarinic treatment [14]. Coincident with the emergence of the abnormal movements in these mice, striatal ChI selectively degenerate, while all other torsinA deficient populations (including other forebrain cholinergic neurons) remain intact. Surviving ChI are hypertrophic and display altered functional connectivity [14] and exhibit gene expression changes consistent with altered synaptogenesis [46].

Here, we explore whether dysfunction of surviving ChI contributes to abnormal movements in Dlx-CKO mice. To directly test the role of ChI degeneration in motor abnormalities, we generated a novel genetic rescue model in which torsinA expression is selectively restored in ChI, which prevents cell loss and partially rescues motor deficits. This demonstrates a requirement for torsinA in ChI for normal motor function. We find that surviving ChI exhibit increased activity following the onset of degeneration and motor abnormalities in Dlx-CKO mice, suggesting that altered cholinergic signaling emerges following degeneration of neighboring cells. We demonstrate that systemic antimuscarinic treatment during the juvenile critical period for torsinA function produces sustained long-term behavioral improvement. Infusion of antimuscarinics directly to the striatum abolishes abnormal clasping, demonstrating that the striatum is the site of action for this behavioral rescue. However, selective ablation of striatal ChI during either postnatal maturation or adulthood does not change motor function in Dlx-CKO mice, suggesting that removal of aberrant ChI function is not sufficient to rescue behavior. Surprisingly, antimuscarinic treatment retains the ability to rescue abnormal clasping movements even after ChI ablation, showing that the behavioral benefit of muscarinic receptor antagonism does not require the continued presence of ChI signaling. Together, these findings redefine the relationship between ChI degeneration, cholinergic dysfunction, and motor abnormalities in a symptomatic model of DYT-TOR1A.

## Results

### Re-expression of Tor1a in striatal cholinergic interneurons rescues ChI degeneration and reduces clasping behavior

All striatal neurons lack torsinA in Dlx-CKO mice, but only a subset of striatal ChI selectively degenerate [14]. Mice lacking torsinA selectively in cholinergic cells also show selective ChI degeneration, suggesting this process is cell autonomous [47]. To determine if ChI degeneration contributes to motor abnormalities, we generated a novel knock-in mouse with selective torsinA expression in cholinergic neurons (*Chat-Tor1a* mice; Figure 1A). *Chat-Tor1a* mice were born at the expected Mendelian ratio, survived, and grew at the same rate as littermate controls (Figure 1D). TorsinA expression was selectively increased in ChI, but not nearby parvalbumin+ (PV) interneurons (Figure 1B-C). To determine if ChI-specific torsinA re-expression in Dlx-CKO prevents ChI degeneration, we performed unbiased stereology of ChAT-immunostained brains. ChAT+ neurons were significantly reduced at postnatal day 70 (P70) in Dlx-CKO mice and ChI-selective torsinA expression prevented this cell loss (Figure 1E). Limb clasping duration was significantly shorter in Dlx-CKO mice expressing *Chat-Tor1a* (Figure 1F), but locomotor hyperactivity was not affected by the *Chat-Tor1a* allele (Figure 1G).

**Figure 1:**
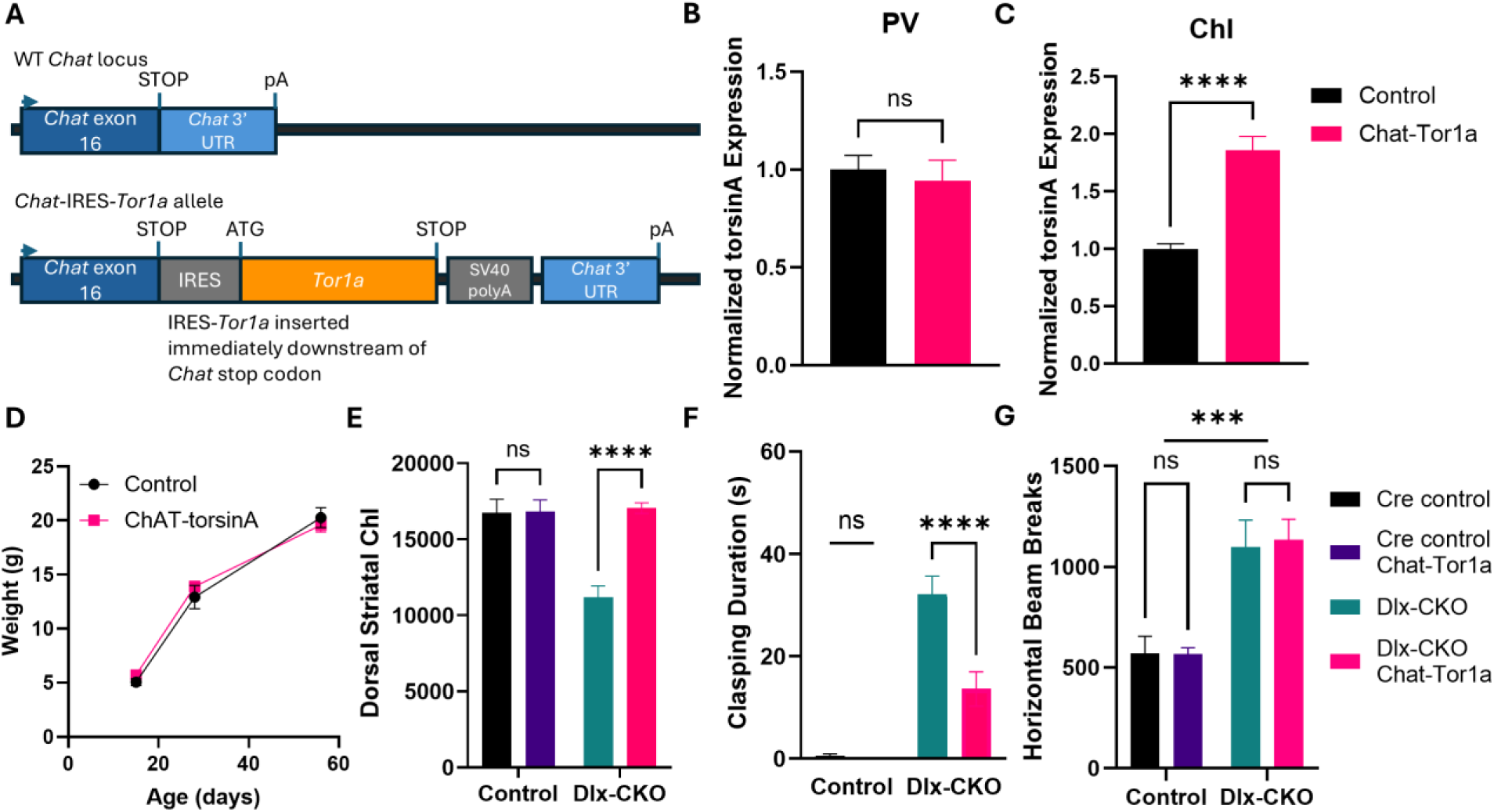
Selective re-expression of *Tor1a* in ChI rescues ChI degeneration and reduces limb clasping duration. (A) Schematic of the ChAT-Tor1a construct (B) Validation of torsinA levels in PV+ interneurons (Welch’s unpaired t test: t=0.4508, df=56.92, p = 0.6539) (C) Validation of torsinA levels in ChI. (Welch’s unpaired t test: t=6.789, df=72.66, p<0.0001) (D) Postnatal weight gain of ChAT-Tor1a mice and Dlx-CKO mice. (Two-way ANOVA: Main effect of genotype F_(1,6)_ = 0.1380, p = 0.723; Main effect of Age F_(2,12)_ = 676.4, p < 0.0001; Age x Genotype F_(2,12)_ = 2.468, p = 0.1266) (E) Quantification of striatal cholinergic interneurons in the ChAT-Tor1a mouse model. (Two-way ANOVA: Main effect of Control vs Dlx Genotype F_(1,28)_ = 13.29, p = 0.0011; Main effect of ChAT-TorA allele F_(1,28)_ = 16.56, p = 0.0003; Interaction F_(1,28)_ = 15.88, p = 0.0004 (F) Clasping duration of ChAT-Tor1a mice at P70. (Two-way ANOVA: Main effect of control vs Dlx Genotype F_(1,54)_ = 57.97, p < 0.001; Main effect of ChAT-TorA allele F_(1,54)_ = 10.35, p = 0.0022; Interaction: F_(1,54)_ = 9.085, p = 0.0039; Šídák’s multiple comparisons test p < 0.0001) (G) Locomotor activity of ChAT-Tor1a mice. (Two-way ANOVA: Main effect of Control vs Dlx Genotype F_(1,31)_ = 35.06, p < 0.0001; Main effect of ChAT-TorA allele F_(1, 31)_ = 0.03073, p = 0.8620; Interaction F_(1,31)_ = 0.04569, p = 0.8321)

### Dlx-CKO dorsal striatal ChI display altered activity beginning during a critical period

Surviving ChI in adult Dlx-CKO mice exhibit abnormal morphological and electrophysiological characteristics following degeneration of neighboring cells [14], suggesting that aberrant signaling of remaining ChI could contribute to abnormal movements. The phosphorylation of ribosomal protein s6 (pRPS6) is correlated with the physiological activity of striatal ChI [48]. To determine how striatal ChI activity changes relative to neurodegeneration and motor dysfunction in Dlx-CKO mice, we quantified pRPS6 fluorescence intensity at three disease stages: before the onset of neurodegeneration (P11), immediately after the emergence of ChI loss and motor abnormalities (P17), and after neurodegeneration is complete and motor phenotypes are stabilized (P56). Striatal ChI pRPS6 staining intensity did not differ between Dlx-CKO mice and controls prior to the onset of neurodegeneration (Figure 2B), but was significantly increased in Dlx-CKO mice at both P17 and P56 (Figure 2B). ChI pRPS6 intensity was not different between genotypes in the ventral striatum, where ChI do not degenerate in Dlx-CKO mice (Figure 2C).

**Figure 2.**
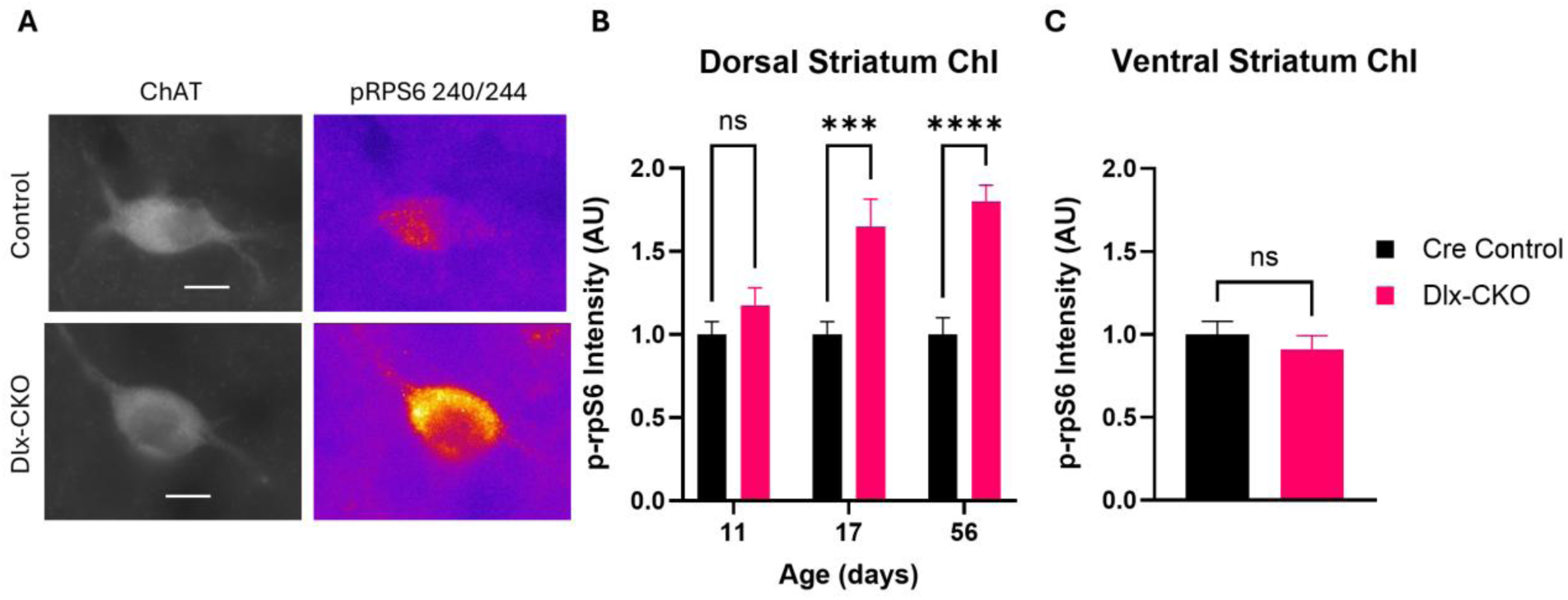
Striatal ChI display increased markers of activity as abnormal clasping movement emerges. (A) Representative images of immunohistochemically stained ChI from the dorsal striatum at P56. Scale bar = 20 μm (B) pRPS6 quantification of dorsal striatal ChI. (Two-way ANOVA: Main effect of Age F_(2,27)_ = 4.644, p = 0.0185; Main effect of Genotype F_(1,27)_ = 37.68, p < 0.0001; Interaction F_(2,27)_ = 4.644; Šídák’s multiple comparisons test: Cre control vs Dlx-CKO: P11 p = 0.2587; P17 p = 0.0004; P56 p < 0.0001) (C) pRPS6 Quantification of ventral striatal ChI. (Welch’s unpaired t test: t=0.7979, df=9.950, p = 0.4435)

### Systemic antimuscarinic administration during striatal maturation reduces limb clasping

We previously reported that acute antimuscarinic administration transiently reduces limb clasping severity in adult mice [14]. However, it is unclear whether earlier antimuscarinic treatment during striatal development would provide longer lasting benefit. Antimuscarinic treatment is more efficacious when instituted early in the disease course, especially in younger patients who are able to tolerate higher doses [49, 50]. Therefore, we treated Dlx-CKO mice beginning at the onset of limb clasping behaviors and continuing through striatal physiological maturation and behavioral stabilization (P14-28). This time range also corresponds to the most active period of striatal ChI degeneration [14]. Dlx-CKO mice were randomly assigned to receive daily intraperitoneal (IP) injections of either 5 mg/kg trihexyphenidyl (THP) or vehicle. 30 minutes following each IP injection, mice underwent a 1-minute tail suspension test (Figure 3A). At baseline and during the first four days of the experiment, when clasping behavior has not yet fully emerged in Dlx-CKO mice [14], both saline- and THP-treated groups exhibited minimal clasping. As the experiment progressed, saline-treated mice developed progressively longer clasping durations, while THP-treated mice showed no increase in clasping (Figure 3A). Following the 14-day treatment period, mice received no additional injections and were monitored into adulthood. The behavioral benefit of juvenile THP treatment persisted long after drug washout, with THP-treated mice continuing to exhibit significantly shorter clasping durations than saline-treated mice. This sustained improvement contrasts with previous studies of adult mice, in which behavioral benefit was not sustained following drug washout [14].

**Figure 3.**
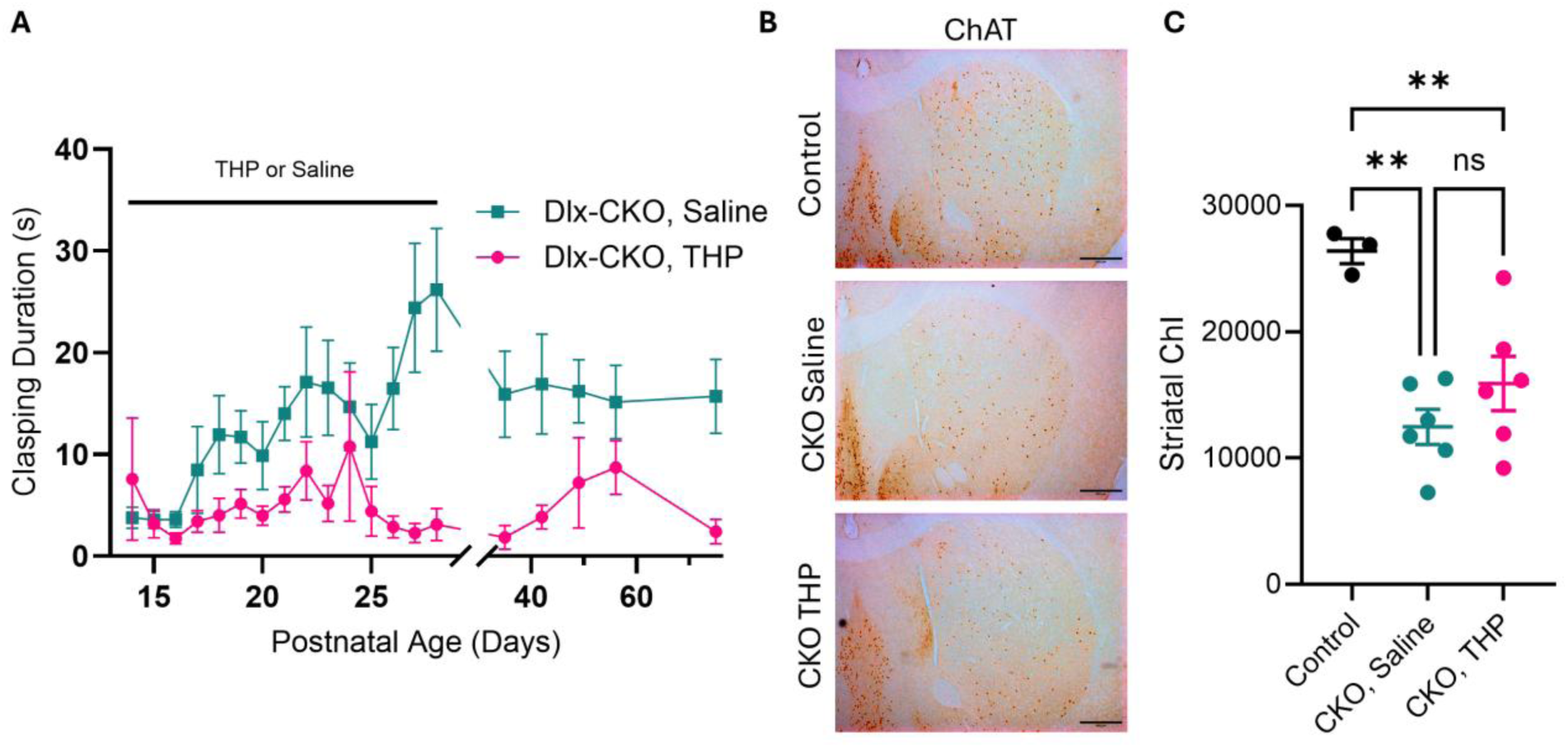
Antimuscarinic treatment during development produces both acute and sustained reductions in abnormal clasping (A) Limb clasping duration of mice treated with the antimuscarinic THP or saline from postnatal day 14-28 (Two-way ANOVA: main effect of Treatment F_(1, 10)_ = 18.20 p = 0.0016, main effect of Time F_(19, 190)_ = 2.086 p = 0.0067, Treatment x Time F_(19, 190)_ = 2.139 p = 0.0052 (B) Representative coronal brain sections stained for ChAT from control, Dlx-CKO mice receiving saline, or Dlx-CKO mice treated with THP during adolescence. Scale bar = 500 μm (C) Quantification of striatal cholinergic interneurons via stereology (Two-way ANOVA F_(2, 12)_ = 11.66 p = 0.0015; Tukey’s multiple comparisons test Control vs CKO, Saline p = 0.0012, Control vs CKO, THP p = 0.0093, CKO, Saline vs CKO, THP p = 0.3471)

In Dlx-CKO mice, ChI degeneration coincides with the emergence of abnormal clasping behavior. Striatal ChI are initially generated and migrate normally but undergo selective degeneration during a critical period of striatal maturation. We previously showed that restoring torsinA expression to all striatal neurons during this developmental window promotes ChI survival and reduces motor abnormalities [23], linking torsinA-dependent ChI pathology to limb clasping. Consistent with this relationship, *Chat-Tor1a* mice exhibited both reduced clasping duration and rescue of ChI degeneration (Figure 1E-F). We next asked whether juvenile THP treatment similarly improved the behavior by preventing ChI loss. Following completion of the behavioral study, we performed unbiased stereological analyses of ChAT-immunostained striatal sections. THP did not increase ChI survival relative to saline, demonstrating that persistent behavioral benefit did not depend on preventing ChI degeneration (Figure 3B-C).

### Juvenile Ablation of ChI does not prevent limb clasping

These findings suggested that genetic torsinA restoration and antimuscarinic treatment improve behavior through distinct mechanisms: torsinA restoration preserves ChI, whereas THP acts despite their continued degeneration. One possibility was that THP instead suppresses aberrant cholinergic signaling originating from the surviving ChI, which exhibited increased pRPS6 during striatal maturation. If these pathological surviving neurons drive motor dysfunction, their removal should phenocopy the behavioral effects of antimuscarinic treatment. To test whether the remaining striatal ChI are required for the emergence of clasping, we infused either a cholinergic-specific saporin toxin (ChAT-SAP, ATS Bio) or saline bilaterally into the striatum of Dlx-CKO mice at P14 (before clasping onset) and performed repeated tail suspension testing into adulthood.

As expected, vehicle-infused Dlx-CKO animals exhibited fewer striatal ChI than cre-control animals at P50, and ChAT-SAP administration significantly reduced the number of remaining striatal ChI in Dlx-CKO mice (Figure 4A-B). To determine the full extent of the lesion, we examined ChAT-immunostained serial coronal sections across the rostral to caudal extent of the striatum. Lesions were confined to the dorsal striatum (caudate-putamen); ChI in the nucleus accumbens, midline septum, and basal forebrain were spared (Figure 4A). Essentially all ChI cell bodies were ablated in rostral sections of the dorsal striatum, while a subset remained in caudal sections containing the tail of the striatum (Figure 4C). Consistent with our previous findings [14], the most severe degeneration in Dlx-CKO mice was observed in the dorsolateral (DL) and dorsomedial (DM) quadrants of the caudate-putamen, where ChAT-SAP treatment further reduced ChI numbers (Figure 4D).

**Figure 4.**
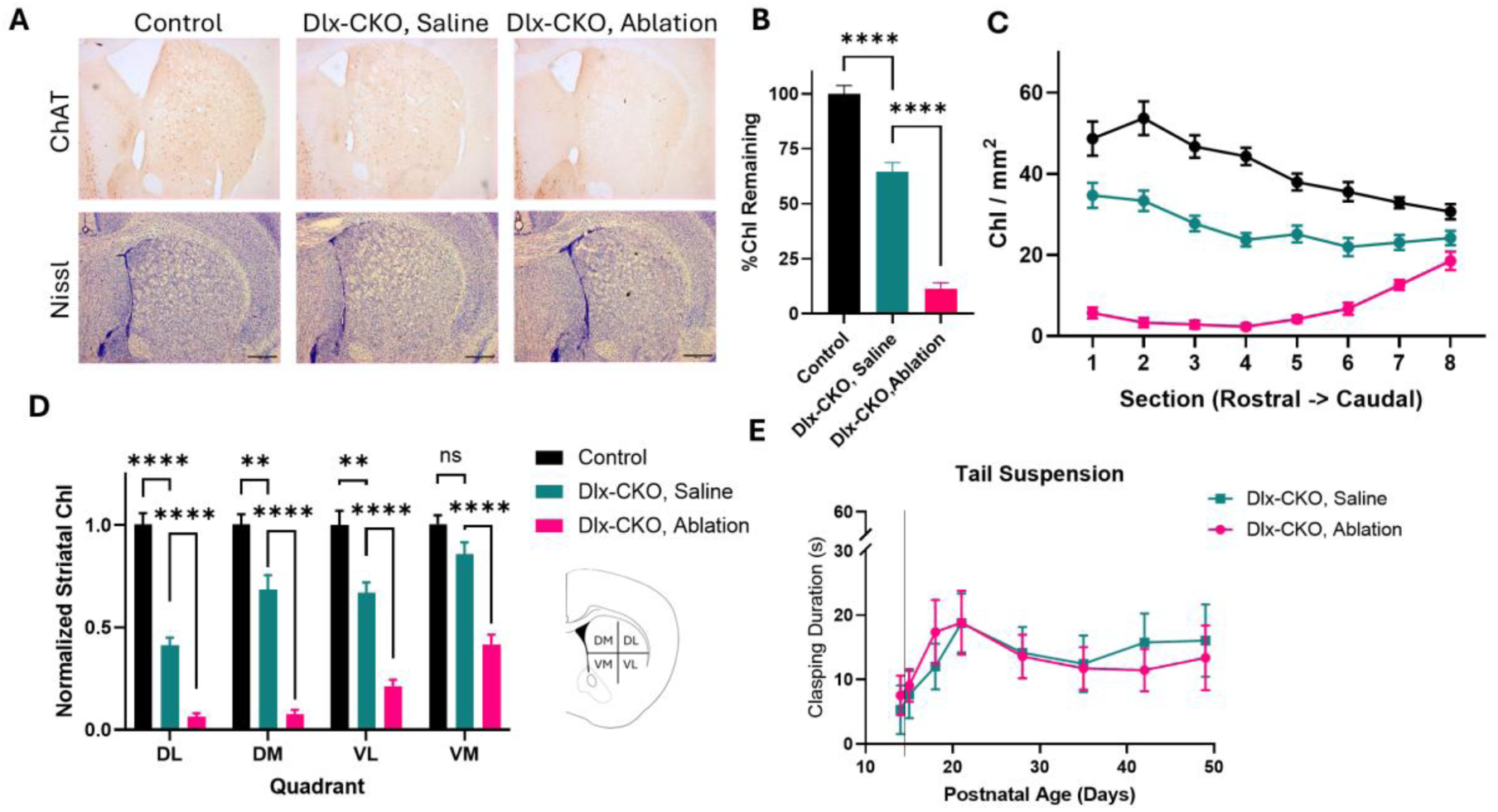
Ablation of striatal ChI at P14 does not affect clasping. (A) Representative histology images from Control, Dlx-CKO with saline infusions into their bilateral caudate-putamen, and Dlx-CKO mice with selective ChI ablation using ChAT-Sap. Top row: Coronal sections stained for ChAT. Bottom Row: Coronal sections stained with Nissl. Scale bar = 500 μm (B) Quantification of dorsal striatal ChI, expressed as % control. Dlx-CKO mice showed a significant decrease in overall striatal ChI number, and ablation of striatal ChI nearly eliminated the remaining ChI. (One-way ANOVA, F_(2,28_) = 149.3, p<0.0001; Tukey’s multiple comparisons: Control vs CKO –Ablation p<0.0001; Control vs CKO +Ablation p<0.0001; CKO –Ablation vs CKO +Ablation p<0.0001) (C) Quantification of striatal ChI by section from rostral to caudal. (Two-way ANOVA, main effect of section F_(3.365, 96.63)_ = 15.81, p<0.0001; Main effect of Genotype F_(2,30)_ = 115.7, p<0.0001; Section x Genotype F_(6.731, 96.63)_ = 21.00, p<0.0001) (D) Quantification of remaining neurons by quadrant, normalized to control (Two-way ANOVA: Main Effect of ChI Status F_(6, 87)_ = 10.62, p <0.0001; Main Effect of Quadrant F_(1.896, 54.99)_ = 33.41, p <0.0001). (E) Striatal ChI ablation does not prevent clasping development (Two-way ANOVA: Main Effect of Treatment F_(1, 22)_ = 0.01658, p = 0.8987; Main Effect of Time F_(6,132)_ = 3.690, p=0.0020; Treatment x Time F_(6,132)_ = 0.4818, p=0.8213). Vertical line denotes time of ablation surgery.

The onset, progression, and duration of limb clasping during tail suspension were not significantly different between animals that received saline and those that received ChAT-SAP (Figure 4E).

### Ablation of striatal cholinergic interneurons does not alter limb clasping behavior in adult mice

Having found that juvenile ChI ablation did not prevent the emergence of clasping, we next asked whether surviving ChI were required to maintain established motor abnormalities in adulthood. Surviving ChI in Dlx-CKO mice exhibit persistent aberrant activity (Figure 2;[14]), suggesting that dysfunction of the remaining cells may contribute to ongoing abnormal motor activity after neurodegeneration. To determine whether ChI ablation alters established clasping behavior after striatal maturation, mice were randomly assigned to receive bilateral striatal infusions of ChAT-SAP or vehicle. ChAT-SAP significantly reduced striatal ChI numbers (Figure 5A-B), especially in the most rostral sections, with the most striking reduction occurring within the ventral quadrants of the dorsal striatum (Figure 5C-D).

**Figure 5.**
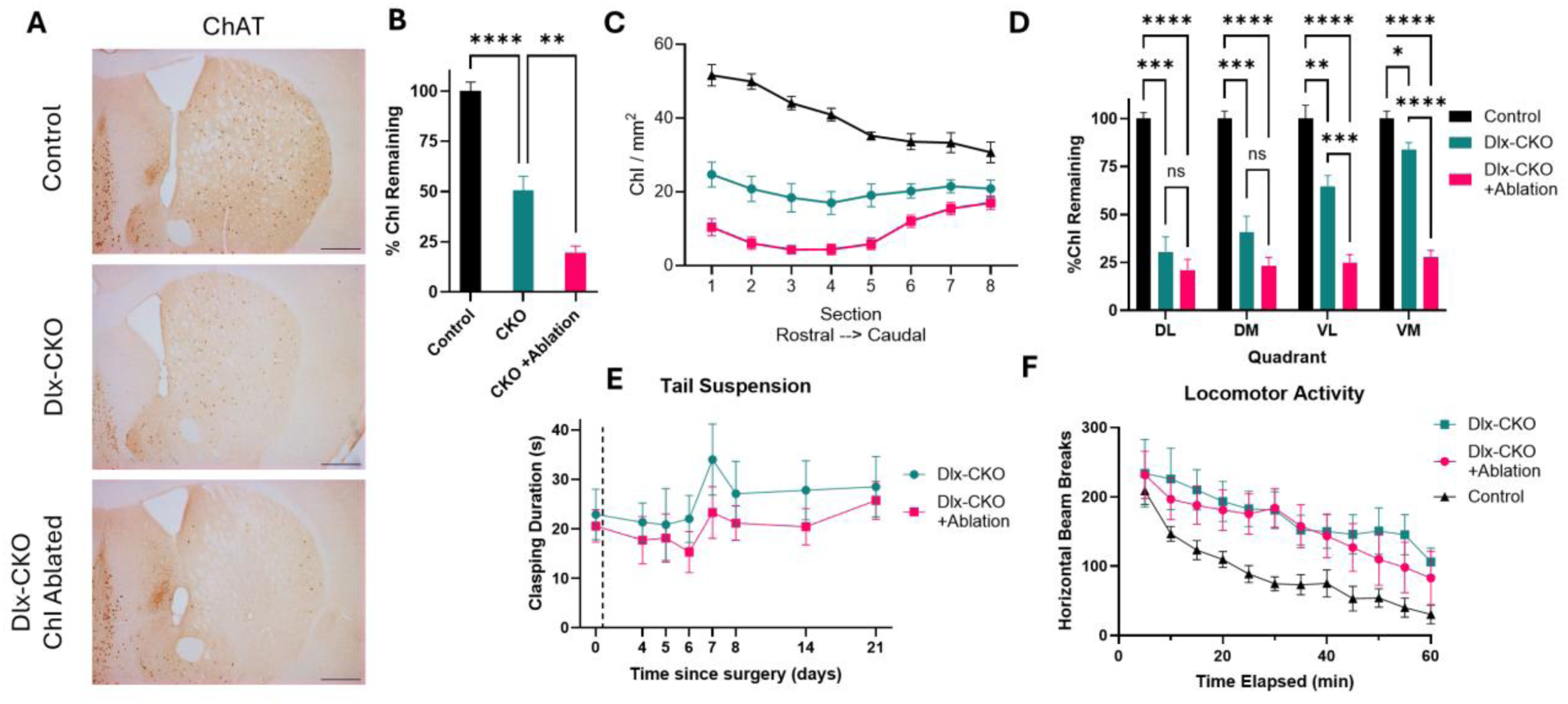
Ablation of striatal ChI at >P70 does not affect clasping duration or hyperactivity. (A) Representative images of dorsal striatum histochemical staining for ChAT. Scale bar = 500 μm (B) Quantification of ChI within the dorsal striatum normalized to control. Dlx-CKO mice have significantly decreased number of striatal ChI than littermate controls. Selective ablation of ChI results in a further significant decrease in ChI. One-way ANOVA, F_(2, 15)_ = 61.5, P<0.0001; Tukey’s multiple comparisons test: Control vs CKO –Ablation p<0.0001; Control vs CKO +Ablation p<0.0001; CKO-Ablation vs CKO+Ablation p = 0.002) (C) Quantification of ChI counted within the dorsal striatum by serial section. Sections are from approximately AP +1.25 mm to –0.5 mm. (Two way ANOVA: Main effect of Section F_(3.535, 52.51_) = 13.65, p< 0.0001; Main effect of ChI Status F_(2, 15)_ = 62.51, p<0.0001, Interaction F_(7.069, 52.51)_ = 19.04, p<0.0001) (D) Quantification of ChI counted within the dorsal striatum by quadrant. (Two-way ANOVA: Main effect of quadrant F_(2.187, 32.8)_ =17.29, p<0.0001; Main effect of Genotype F_(2,15)_ = 55.99, p<0.0001; Quadrant x Genotype Interaction F_(4.373, 32.8_0) = 17.35, p < 0.0001 (E) Ablation of ChI in adult Dlx-CKO mice does not reduce clasping duration (Two-way ANOVA: Main Effect of Treatment F_(1, 10)_ = 0.2260, p = 0.6447. Vertical line denotes time of ablation surgery (F) Ablation of ChI does not alter locomotor hyperactivity. Two-way ANOVA, main effect of genotype F_(2, 19)_ = 2.571, P=0.1027

Despite the significant reduction in striatal ChI number, Dlx-CKO mice with ChI ablation showed no change in the duration of their clasping relative to vehicle-treated animals (Figure 5E). ChI ablation also did not alter the locomotor hyperactivity of Dlx-CKO mice (Figure 5F)[14].

We next sought to confirm that the loss of ChI cell bodies was accompanied by a reduction in striatal cholinergic machinery. ChI produce and release the majority of acetylcholine in the striatum [51, 52] although additional cholinergic axon terminals originate from the pedunculopontine nucleus [53, 54]. Striatal ChI also express high levels of acetylcholinesterase (AChE), which rapidly terminates cholinergic signaling [55–57]. To assess the impact of ChI ablation on AChE activity, unilaterally infused ChAT-SAP into the striatum of adult Dlx-CKO mice and assessed AChE activity two weeks later. Consistent with our previous findings [14], Dlx-CKO mice exhibited significantly less AChE activity than littermate controls in the unablated dorsal striatum (Figure 5S). Within Dlx-CKO mice, AChE staining was significantly reduced in the ablated hemisphere relative to the unablated hemisphere, confirming that ChI ablation substantially reduced striatal AChE activity (Figure 5S).

### Intrastriatal antimuscarinic administration reduces limb clasping

The antimuscarinic trihexyphenidyl is effective in reducing dystonia symptoms at high doses [13, 58]. Muscarinic receptor subtypes are expressed widely in the brain, including in the striatum, cortex, basal forebrain, globus pallidus, substantia nigra, midbrain tegmentum, and cranial nerve nuclei [59, 60], making the site of its therapeutic effects unclear. We hypothesized that striatum is the principal site at which muscarinic antagonism reduces clasping in Dlx-CKO mice.

To restrict muscarinic antagonism to the striatum, we bilaterally implanted Dlx-CKO mice with chronic infusion cannulae targeting the dorsal striatum (Figure 6A-B). Due to solubility limitations of trihexyphenidyl in aCSF, we infused scopolamine, another nonselective antimuscarinic agent that we previously demonstrated significantly reduces clasping in Dlx-CKO mice [14]. Thus, this experiment tested whether muscarinic antagonism within the striatum is sufficient to reduce clasping. Mice underwent a 1 minute tail suspension before and 30 minutes after bilateral infusion of scopolamine or aCSF to the striatum. Both low (5 µg per hemisphere) and high (30 µg per hemisphere) dose scopolamine significantly reduced clasping duration compared to vehicle, with the high dose nearly eliminating clasping behaviors in most animals (Figure 6C, 6S).

**Figure 6.**
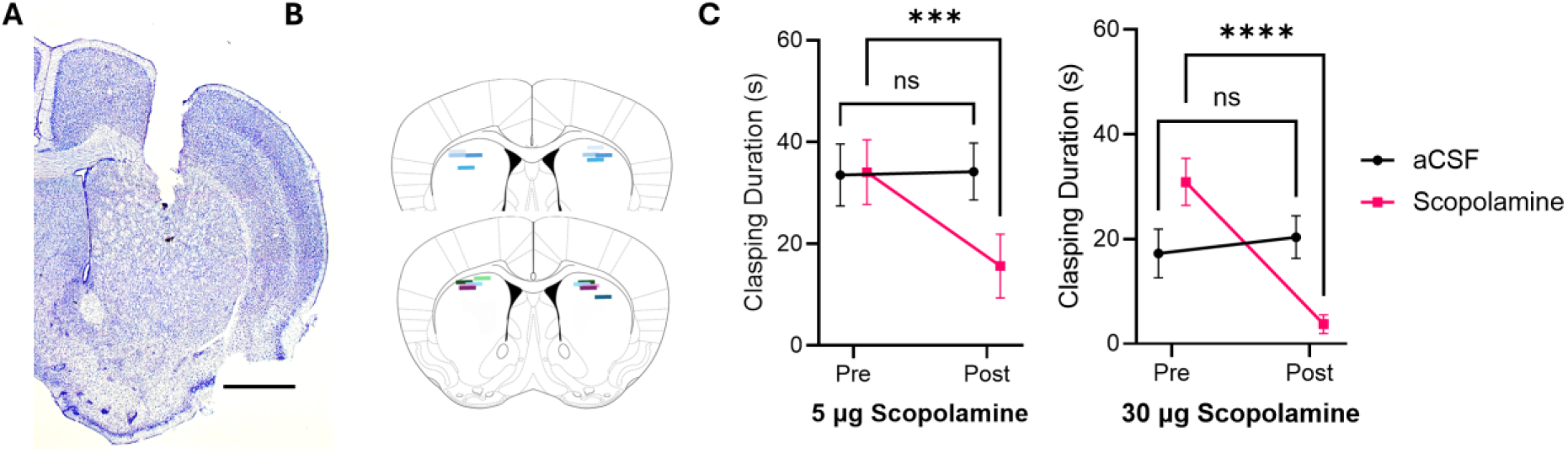
Intrastriatal antimuscarinic treatment significantly reduces limb clasping duration in Dlx-CKO mice (A) An example Nissl-stained section of a cannulated animal, demonstrating the cannula termination within the dorsal striatum. Scale bar = 500 μm (B) Schematic of coronal brain slices showing cannula placement within the dorsal striatum. Horizontal lines depict the location of the end of the cannula as determined by histology. Each color represents a single animal. Coronal sections corresponding to AP +0.7 and +0.8 mm from bregma (C) Limb clasping durations pre and post infusion of either aCSF or Scopolamine bilaterally. Scopolamine significantly reduced clasping duration following both low (5 μg) and high (30 μg) dose infusions. (Two-way ANOVA; 5 ug scopolamine Time x Treatment F_(1,9)_ = 12.43, p=0.0065); 30 ug scopolamine Time x Treatment F_(1,9)_ = 46.64, p <0.0001)

### Antimuscarinics reduce limb clasping in ChI-ablated Dlx-CKO mice

Having established that muscarinic antagonism within the striatum is sufficient to reduce clasping, we next asked whether surviving ChI are required for this effect. All five muscarinic subtypes are expressed within the striatum with significant heterogeneity in cellular and subcellular distributions [37, 61, 62]. Inputs to the striatum, including those arising from the cortex, thalamus, and SNc differentially express M2/4, M3, and/or M5 receptors presynaptically [34, 63–66]. Both D1R-expressing and D2R-expressing SPNs express M1 receptors, whereas D1R neurons also express M4 receptors [63, 65, 67]. ChI predominantly express M2/M4 receptors, which function as inhibitory autoreceptors [63, 67–70]. Given the increased ChI activity of surviving ChI (Figure 2), and established alterations in ChI excitability [14], we hypothesized that muscarinic receptors associated with the surviving ChI might contribute to the behavioral effects of antimuscarinic treatment.

To test this hypothesis and determine whether surviving striatal ChI are required for the antimuscarinic response, we performed bilateral infusions of ChAT-SAP or vehicle into the dorsal striata of adult Dlx-CKO mice or control mice. As in previous experiments, unablated Dlx-CKO mice had significantly fewer striatal ChI than controls, and ablated Dlx-CKO mice showed a further significant decrease in overall ChI number (Figure 7A). ChAT-SAP administration significantly reduced ChI throughout the rostral-caudal extent of the striatum, but did not fully eliminate all ChI, particularly in caudal sections (Figure 7B). ChI reductions were similar across all quadrants of the dorsal striatum compared to unablated Dlx-CKO mice and littermate controls (Figure 7C). As with previous ablation experiments, ChI loss was limited to the caudate-putamen, and did not affect cholinergic neurons in the basal forebrain, medial septum, or nucleus accumbens (data not shown). Importantly, the magnitude and pattern of ChI loss was equivalent between animals that received THP and those that received saline (Figure 7A-C). One week following surgery, mice received daily IP injections of THP or saline for five days and underwent tail suspension testing 30 minutes following each daily injection. Both ChI-ablated and nonablated Dlx-CKO mice showed a significant reduction in clasping duration following systemic THP treatment relative to their respective saline-treated controls (Figure 7D). ChI-ablated and nonablated miceshowed similar reduction in clasping duration during THP treatment, indicating that extensive depletion of dorsal striatal ChI did not diminish the systemic antimuscarinic response.

**Figure 7.**
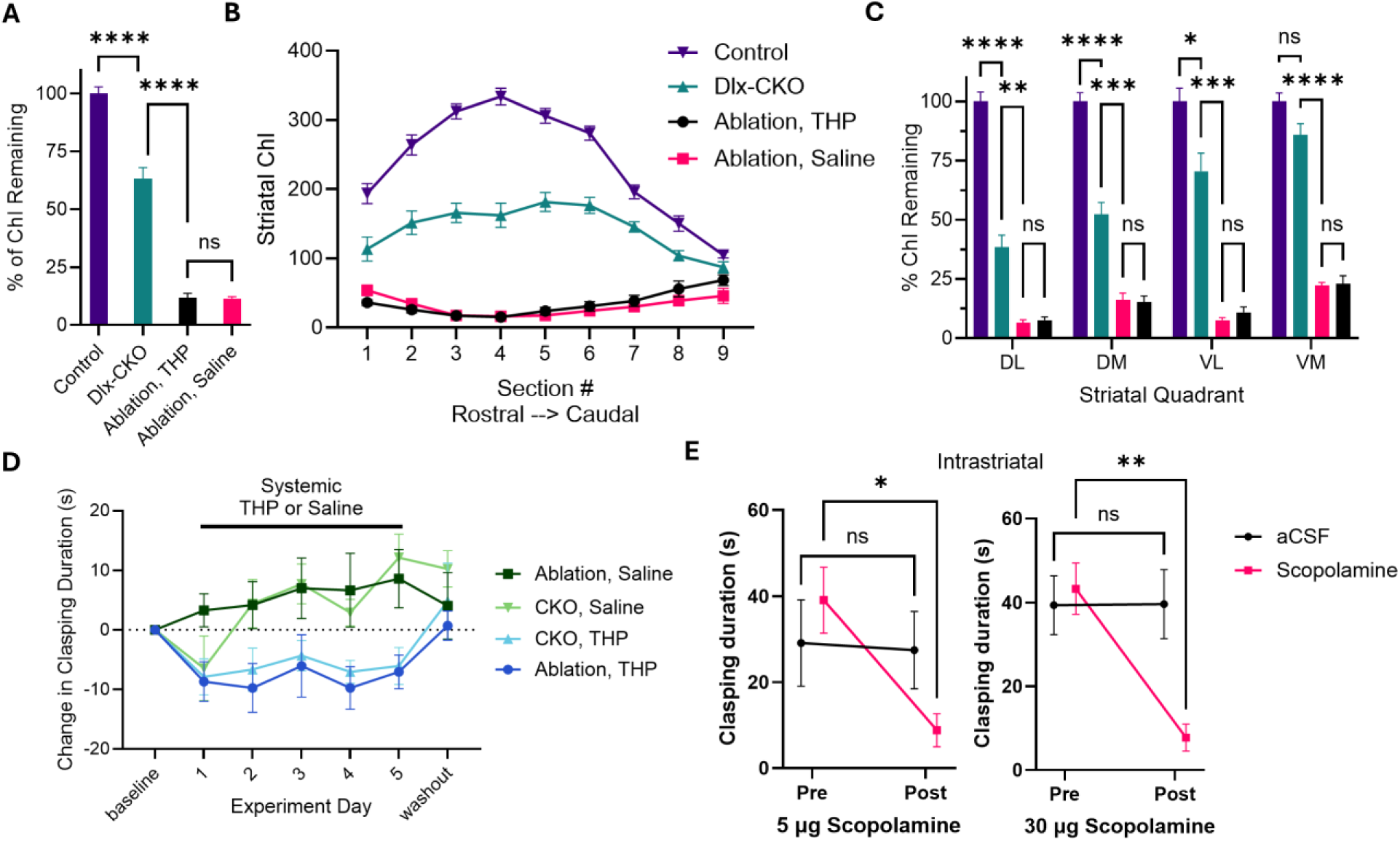
Both systemic and intrastriatal antimuscarinics reduce clasping in Dlx-CKO mice with striatal ChI ablations. (A) Quantification of striatal ChI, expressed as % of control. One-way ANOVA; Main effect of ChI status F_(3,32)_ = 275.8, p<0.0001; Tukey’s multiple comparisons: Control vs CKO – Ablation p<0.0001; CKO-Ablation vs CKO +Ablation Saline p<0.0001; CKO –Ablation vs CKO +Ablation THP p<0.0001; CKO +Ablation Saline vs CKO +Ablation THP p = 0.9991) (B) Quantification of striatal ChI from the rostral to caudal extent of the striatum (Two-way ANOVA: Main effect of ChI Status F_(3, 32)_ = 278.5, p <0.0001; Main effect of Section F_(3.214, 102.8)_ = 20.39, p < 0.0001; Section x ChI Status F_(21, 224)_ = 28.04, p < 0.0001) (C) Remaining ChI per striatal quadrant, expressed as % remaining of control (Two-way ANOVA, Quadrant x ChI status F_(9, 96)_ = 10.49, p <0.0001) (D) Limb clasping duration for ablated and unablated Dlx-CKOs given IP THP or vehicle; data are presented as the change in clasping duration from baseline (Two-way ANOVA, main effect of group F_(3, 24)_ = 3.560 p = 0.0292; Time x Group F_(18, 144)_ = 2.129 p = 0.0075). (E) Scopolamine infusion reduces clasping in Dlx-CKO mice with striatal ChI ablations (Two-way ANOVA; 5 ug Pre/post x Drug F_(1,5)_ = 16.49, p=0.0097; 30 ug Pre/Post x Drug F_(1, 5)_ = 50.73, p=0.0008)

Because systemic THP could act at muscarinic receptors outside the striatum, we next asked whether intrastriatal antimuscarinic treatment remained effective after ChI ablation. Dlx-CKO mice received bilateral intrastriatal ChAT-SAP administration followed by bilateral implantation of infusion cannulae targeting the dorsal striatum. Mice were randomly assigned to receive either vehicle (aCSF) or 5 ug of scopolamine per hemisphere on the first experimental day, followed by crossover to receive the other compound after a four-day washout. One week later, the experiment was repeated in the same cohort with a higher dose of scopolamine (30 ug per hemisphere). Both the low- and high-dose of scopolamine significantly reduced clasping duration when delivered directly to the striatal ChI-depleted striatum (Figure 7E, 7S). Thus, extensive depletion of dorsal striatal ChI did not diminish the behavioral efficacy of either systemic or intrastriatal antimuscarinic treatment.

## Discussion

Despite a longstanding association between cholinergic dysfunction and dystonia, neither the causal contribution of ChI dysfunction for the motor phenotype of a DYT1 model nor the anatomic site of therapeutic antimuscarinic action has been established. Our work demonstrates that torsinA-dependent ChI dysfunction causally contributes to the full motor phenotype and that the striatum is the site of therapeutic antimuscarinic action. Selective restoration of torsinA in cholinergic interneurons (ChI) prevented their degeneration and reduced abnormal movements, establishing a causal contribution of torsinA-related ChI dysfunction to motor dysfunction. Antimuscarinics infused directly into the striatum essentially abolished abnormal movements, and systemic treatment during striatal maturation produced benefit that persisted well beyond the treatment period. Together, these findings establish torsinA-dependent ChI dysfunction as a contributor to the motor phenotype and striatal muscarinic signaling as a critical substrate of its pharmacological treatment.

Remarkably, however, the therapeutic benefit of antimuscarinics did not require the continued presence of most dorsal ChIs, dissociating the cellular cause of the motor phenotype from the target of its treatment. Ablation of striatal ChI did not reduce or prevent motor dysfunction, and antimuscarinics remained efficacious even after extensive depletion. Because both the phenotype and its pharmacological suppression persist after extensive depletion of dosrsal striatal ChI, these data support a model in which torsinA-dependent ChI dysfunction disrupts striatal development, producing motor abnormalities that do not require ongoing signaling from most dorsal striatal ChI. As one of the earliest neurons to populate the striatum, ChI are positioned to organize striatal circuit assembly. Their torsinA-dependent dysfunction during this period, when surviving ChI display increased activity and receive abnormal synaptic input, likely establishes an aberrant circuit whose motor consequences persist after most dorsal ChIs are no longer required [31, 32, 71, 72]. These unexpected findings shift a central question in dystonia pathogenesis from how ongoing ChI dysfunction perturbs the mature striatal circuit to how torsinA-dependent ChI pathology disrupts circuit assembly during development, producing motor consequences that persist after extensive depletion of mature dorsal striatal ChI. Separately, the persistence of antimuscarinic efficacy after ChI depletion further dissociates the contribution of ChI dysfunction to the motor phenotype from the mechanism of its pharmacological suppression.

The aberrant striatal circuit is unlikely to arise from ChI dysfunction alone. Because the Dlx-CKO model lacks torsinA in all striatal neurons, the incomplete rescue by ChI-restricted torsinA restoration suggests additional torsinA-dependent contributions from non-ChI striatal cell types. The persistence of motor dysfunction after ChI ablation is also consistent with a circuit abnormality that, once established, no longer requires most dorsal striatal ChI. Together, these experiments implicate torsinA-dependent ChI dysfunction during a striatal maturation in the development of the motor phenotype and identify striatal muscarinic signaling, which remains drug-responsive after extensive ChI depletion, as the critical substrate of antimuscarinic benefit. This framework reconciles the causal contribution of torsinA-dependent ChI pathology with the dispensability of most mature dorsal striatal ChI for both expression of the phenotype and its pharmacological suppression.

The increased activity we observe in surviving ChI, inferred from the activity-dependent marker pRPS6 [48], emerges as abnormal movements appear and is sustained into adulthood. This is consistent with recent work showing that increasing ChI activity induces abnormal movements in mice [22] and that intra-putaminal muscarinic agonists elicit dystonic movements in non-human primates [21]. However, ablating these abnormally active ChI neither prevented nor reduced clasping, and did not alter the response to antimuscarinics, indicating that the behavioral benefit of trihexyphenidyl and scopolamine does not require autoreceptors on ChI. The increased activity of surviving ChI may therefore reflect the aberrant developmental circuit rather than serve as an ongoing driver of the motor phenotype.

A key unresolved question raised by this work is the cellular basis of the muscarinic signaling that is antagonized by antimuscarinics. One possibility is that muscarinic receptors, like many GPCRs, have constitutive activity in the absence of ligand, as observed for the human M3 receptor [73, 74]. If antimuscarinics act as inverse agonists, they could suppress this ligand-independent signaling. Moreover, abnormal striatal development driven by ChI dysfunction may cause misexpression of muscarinic receptors, placing them on striatal cell types that do not normally express them or altering their expression levels. Despite extensive ChI depletion in the dorsal striatum, a small number of cells remained, particularly in the tail of the striatum and in the ventral striatum. The few remaining intrinsic ChI could be sufficient to activate muscarinic signaling cascades, particularly if receptor upregulation occurred in response to ablation [73, 75, 76]. Indeed, ChI have extensive arbors covering large areas of the rodent striatum and influence activity far from their soma [77, 78]. However, the significant reduction in AchE activity observed following ChI ablation confirms a substantial loss of ChI-associated cholinergic machinery, although AChE activity does not directly measure acetylcholine release or signaling [55–57]. Alternatively, extrinsic cholinergic innervation from the pedunculopontine nucleus may contribute the acetylcholine signal antagonized by antimuscarinics [53].

Defining which muscarinic receptors mediate this benefit requires considering their distribution across striatal cell types. All five subtypes are expressed in the striatum, but M1 and M4 predominate [61]. Direct-pathway SPNs express both M1 and M4 receptors, indirect-pathway SPNs express M1, and ChI and some GABAergic interneurons express M2/M4 receptors; on ChI these receptors function as inhibitory autoreceptors [61, 67, 79]. Muscarinic signaling shapes corticostriatal plasticity through these receptors: M1 activation on SPNs is required for LTP induction, M2 and M4 receptors negatively modulate it, and reduced M1 tone permits LTD [66, 80, 81]. Because antimuscarinic benefit persists after extensive dorsal striatal ChI depletion, the relevant receptors are unlikely to reside exclusively on ChI and likely receptors on SPNs or other striatal targets.

The receptor subtypes responsible for the therapeutic effect remain undefined. Trihexyphenidyl and scopolamine are nonselective muscarinic antagonists [58, 82], and the high homology among subtypes has made it difficult to isolate individual contributions in dystonia. Newer compounds with improved M1 or M4 selectivity now make this tractable [83–85]. Ex vivo, selective M1 antagonism rescues SPN plasticity in the Tor1a^+/ΔE^ model [86], M4 antagonism normalizes striatal dopamine release [87], and a selective M4 antagonist is behaviorally efficacious in a dopa-responsive dystonia model [83]. Whether antagonism of M1, M4, or both is required for benefit in DYT-TOR1A dystonia is unknown, and defining this, together with the cell type on which the relevant receptors reside, is an important next step toward subtype-selective therapy.

These findings carry translational implications. DYT-TOR1A dystonia typically presents in childhood or adolescence [88], and antimuscarinic therapy is most effective when initiated early in the disease course [49, 50]. In our model, antimuscarinic treatment restricted to juvenile striatal maturation produced benefit that persisted long after drug withdrawal, suggesting a critical developmental window during which intervention yields durable improvement. Moreover, because antagonism of striatal muscarinic signaling is sufficient to reduce abnormal twisting, subtype- or circuit-selective antimuscarinics could retain efficacy while sparing the off-target effects that limit current nonselective therapy [89].

Together, our findings suggest that ChI degeneration alone is not sufficient to explain locomotor hyperactivity, limb clasping, or the behavioral response to antimuscarinic treatment, despite their similar temporal progression. Rather, they highlight an earlier developmental role for ChI, in which torsinA-dependent dysfunction during striatal maturation shapes the circuit that ultimately drives motor abnormalities. This work redefines the relationship between ChI degeneration, cholinergic dysfunction, and motor abnormalities in a symptomatic model of DYT-TOR1A dystonia.

## Materials and Methods

### Key Resources Table

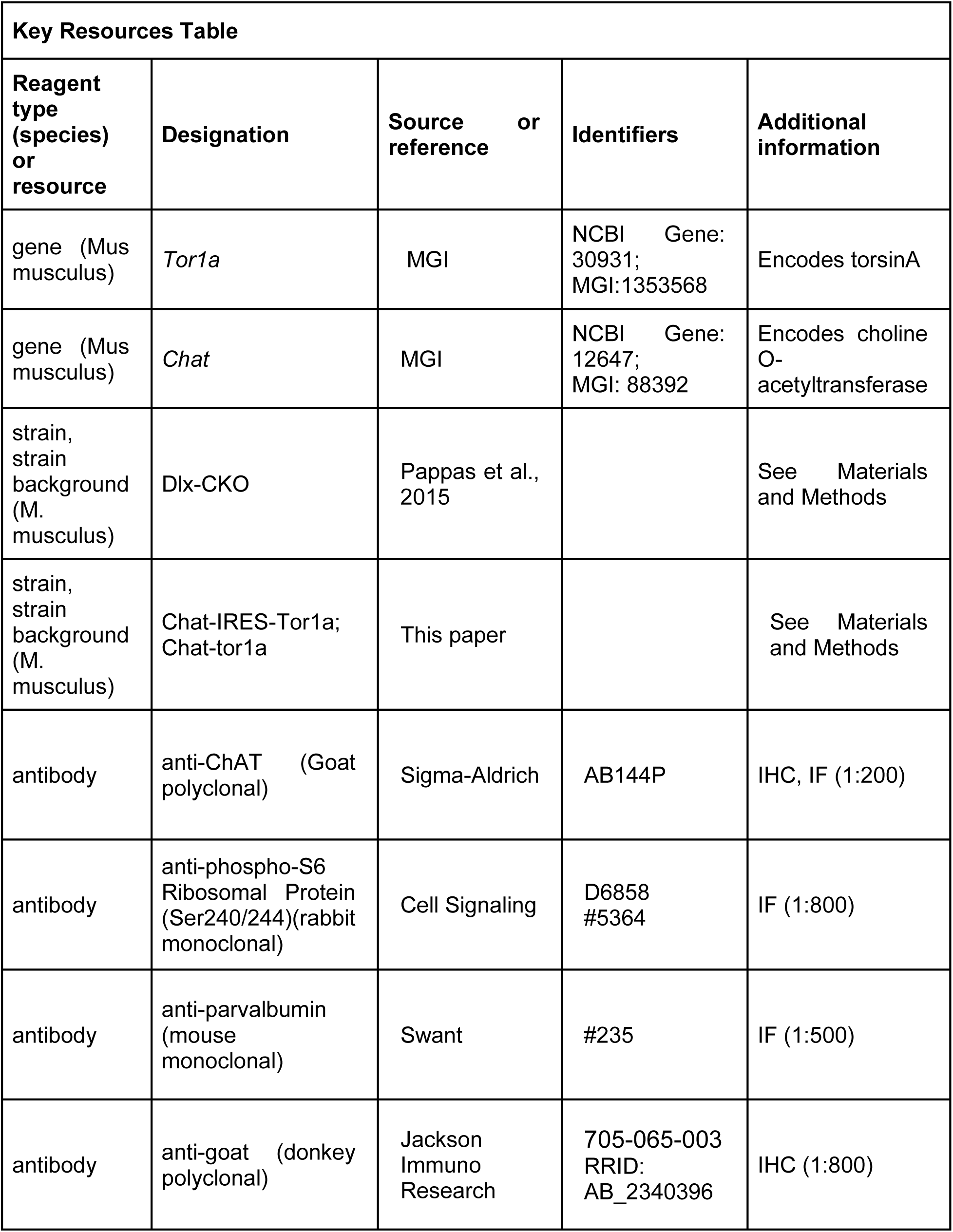
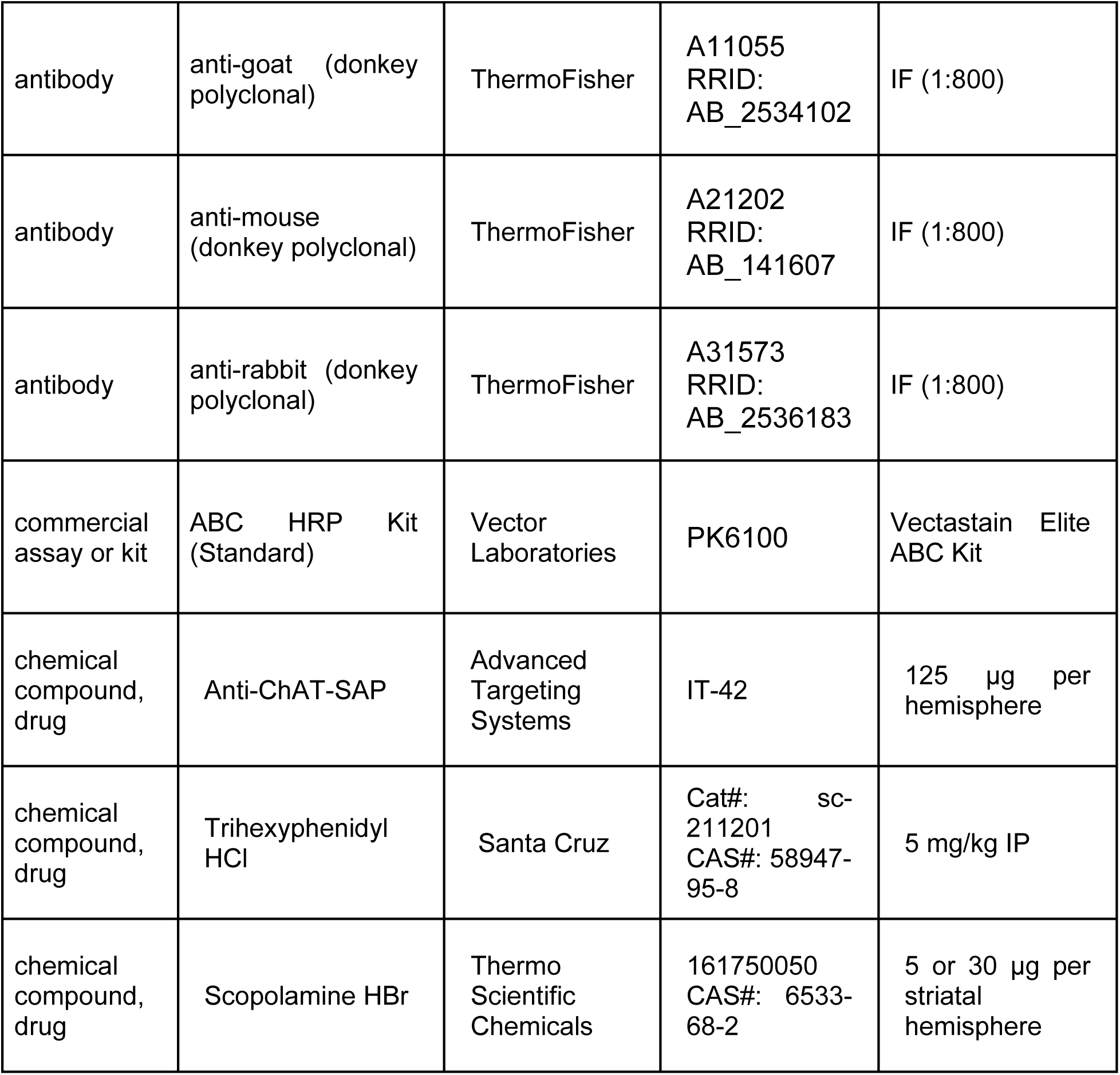

### Animals

Male and female Dlx-CKO mice were used in all studies [Experimental mice: Dlx5/6 cre+, Tor1A^KO/flx^;control mice: Dlx5/6 cre+, Tor1a^flx/+^] [14, 90]. Littermates were used as controls whenever possible; otherwise, age- and sex-matched animals were used. Animals were maintained in 12 hour light/dark cycle (lights on 0600), with food and water provided ad libitum. All behavioral experiments were performed during the light cycle. All animal work has been approved and conducted under the oversight of the UT Southwestern Institutional Animal Care and Use Committee.

*Chat*-IRES-*Tor1a* (*Chat-Tor1a*) knock-in mice were generated using CRISPR/Cas9-mediated genome editing. An sgRNA targeting the region between Chat exon 16 and the 3′ UTR was selected based on in silico design (Wellcome Sanger Institute CRISPR design tool) and validated using a Universal CRISPR Activity Assay (Biocytogen Pharmaceuticals). A circular donor vector containing an IRES-torsinA-SV40 polyA cassette flanked by 1.4-kb left and right homology arms was used for homology-directed repair, resulting in insertion of the IRES-torsinA-polyA sequence immediately downstream of the Chat stop codon. Cas9 mRNA, sgRNA, and donor vector were co-injected into one-cell-stage C57BL/6N embryos, which were subsequently transferred into pseudopregnant Kunming females. Founder mice were identified by PCR and sequencing and crossed to C57BL/6J mice to establish germline transmission. F1 offspring were validated by PCR, Southern blotting, and DNA sequencing.

### Surgery

All surgical procedures were performed under aseptic conditions. Mice were anesthetized with isoflurane in O_2_ and placed into a stereotaxic frame. Analgesia (Buprenorphine SR 1 mg/kg and Carprofen 5 mg/kg) was given subcutaneously at the time of surgery and for 2 days following the procedure. Post-surgery, animals were monitored during recovery on a heating pad before being returned to a clean cage.

Cholinergic Ablation: 250 nl of 0.5 mg/mL IT-42 Anti ChAT-SAP (Advanced Targeting Systems) was administered bilaterally at the injection coordinates (from bregma) AP: +0.6 ML: +/− 2.1 DV: −3.3 mm

Cannula Implantation: Bilateral infusion cannulas (Plastics One) were directed to the dorsal striatum using coordinates (from bregma) AP: +0.5 mm, ML +/− 1.7, DV: −1.6 mm from the dura, and secured to the skull using Metabond. Dummy cannulas were placed to prevent clogging or infection.

### Behavior

Tail suspension: Mice were manually suspended by their tail for 1 minute video recordings and the duration of their forelimb and hindlimb clasping assessed by a minimum of two blinded reviewers for each experiment. In the case of drug administration, suspension was performed 30 minutes following drug administration.

Locomotor Activity: Mice were placed into a fresh cage with new bedding and their horizontal movement and rears were measured via infrared beams (PAS Instruments). Data are presented as activity per 5-minute bins.

Intrastriatal infusion experiments: Mice recovered for a minimum of 3 days following cannula implantation, and a minimum of 3 days elapsed between each infusion day. Animals were removed from their home cages and placed into a clean cage in which they could freely ambulate for the duration of infusions. Scopolamine HBr was delivered bilaterally in a volume of 0.5 or 1 µl at a rate of 200 nl/min via polyethylene tubing attached to a 10 µl Hamilton syringe. The infusion cannula was left in place for an additional 2 minutes to prevent backflow, after which the dummy cannula was replaced and the mouse returned to its home cage. Infusion progress was monitored by observing a small bubble within the tubing advance from its starting position pre-infusion. Animals that did not have confirmed infusions were excluded from analysis.

Juvenile Antimuscarinic: Mice were randomly assigned to experimental groups. Each day from postnatal day 14 to 28, the mice were weighed and received an IP injection of either THP (5 mg/kg, 10 mL/kg volume) or vehicle. 30 minutes post injection the mice were tail suspended. Post-weaning, mice no longer received injections but were tail suspended weekly.

### Compounds

Trihexyphenidyl HCl was obtained from Santa Cruz. It was dissolved in sterile saline and administered IP at a dose of 5 mg/kg. Vehicle was administered at an equivalent volume of 10 mL/kg.

Scopolamine HBr was obtained from Thermo Scientific Chemicals and was dissolved in aCSF (in mM: 92 NaCl, 2.5 KCl, 10 MgSO4, 1 CaCl2, 1.2 NaH2PO4, 30 NaHCO3, 20 HEPES).

### Histology

Mice were deeply anesthetized with a lethal dose of ketamine/xylazine followed by transcardial perfusion with Phosphate Buffered Saline (PBS) followed by 4% paraformaldehyde (PFA) in 0.1M Phosphate Buffer (PB). Brains were removed, post-fixed in 4% PFA for 24 hours, cryoprotected in 20% sucrose, then sectioned at 40 µm on a cryostat (Leica), and stored in PBS with 0.1% sodium azide.

Immunohistochemistry: Free-floating sections were washed with PBS + 0.1% TritonX-100 (PBS-Tx), followed by 20 minutes in 0.3% H_2_O_2_ in PBS. The sections were subsequently washed with PBS, then blocked for 1 hour at room temperature in PBS-Tx + 5% Normal Donkey Serum. The sections were incubated in primary antibody overnight at 4°C. The next day, the sections were washed, then incubated with biotinylated secondary antibody for 1 hour at room temperature. Next, they were incubated with avidin-biotin-peroxidase complex (Vectastain Elite ABC Kit Standard; PK6100, Vector Laboratories) for 1 hour followed by DAB reaction (SigmaFast DAB or SigmaFast DAB with Metal Enhancer). The sections were then mounted onto gelatin-coated slides, dried overnight, dehydrated with a series of ethanol and xylenes, and coverslipped with Permount. Slides were imaged using brightfield microscopy.

Acetylcholinesterase activity: Animals were deeply anesthetized with isoflurane and sacrificed via cervical dislocation. The brain was rapidly removed, frozen on dry ice, and stored at –80°C until cryosectioning. 25 µm serial sections were taken from each brain and stored at –80°C on gelatin coated slides. Cholinesterase histochemistry was performed as previously described [14]. Optical density was determined using ImageJ, with staining from the anterior commissure used for background determination.

### ChI Quantification

ChAT stereology: Striatal ChAT+ neurons were counted using the Optical Fractionator method (MBF Biosciences) using brightfield microscopy. 8 serial coronal brain sections were collected with a section interval of 6. A counting frame of 100×100 µm, grid size of 300×300 µm, guard zone of 1 µm, and 12 µm section thickness was used. Gunderson CE was less than 0.1 for all brains assessed. Boundaries of the caudate putamen (CPu) borders were determined via a reference atlas [91].

Quadrant Analysis and Ablation Quantifications: Serial coronal sections of the striatum (AP: +1.3 mm to –0.9 mm) were stained for ChAT and imaged using brightfield microscopy. The dorsal striatum was divided into quadrants (horizontally between the corpus callosum and anterior commissure; vertically at midpoint of corpus callosum). ChAT+ neurons were manually counted using ImageJ.

### Statistics

All statistical tests were performed using GraphPad Prism 10 or 11. All data are presented as mean ± SEM unless otherwise stated. Animals of both sexes were used; sex was not analyzed as an independent variable. Experimental group sizes were determined using effect sizes of prior behavioral and histological experiments [14, 23]. All animals were randomly assigned to experimental groups. For crossover studies, mice were randomized to receive either treatment or vehicle first; groups were shuffled between drug dose changes. Experiments were repeated at least once to ensure reproducibility.

Data are presented as mean ± SEM unless otherwise specified

*p < 0.05, **p <0.01, ***p <0.001, ****p < 0.0001

---

**Figure 5 Supplemental:**
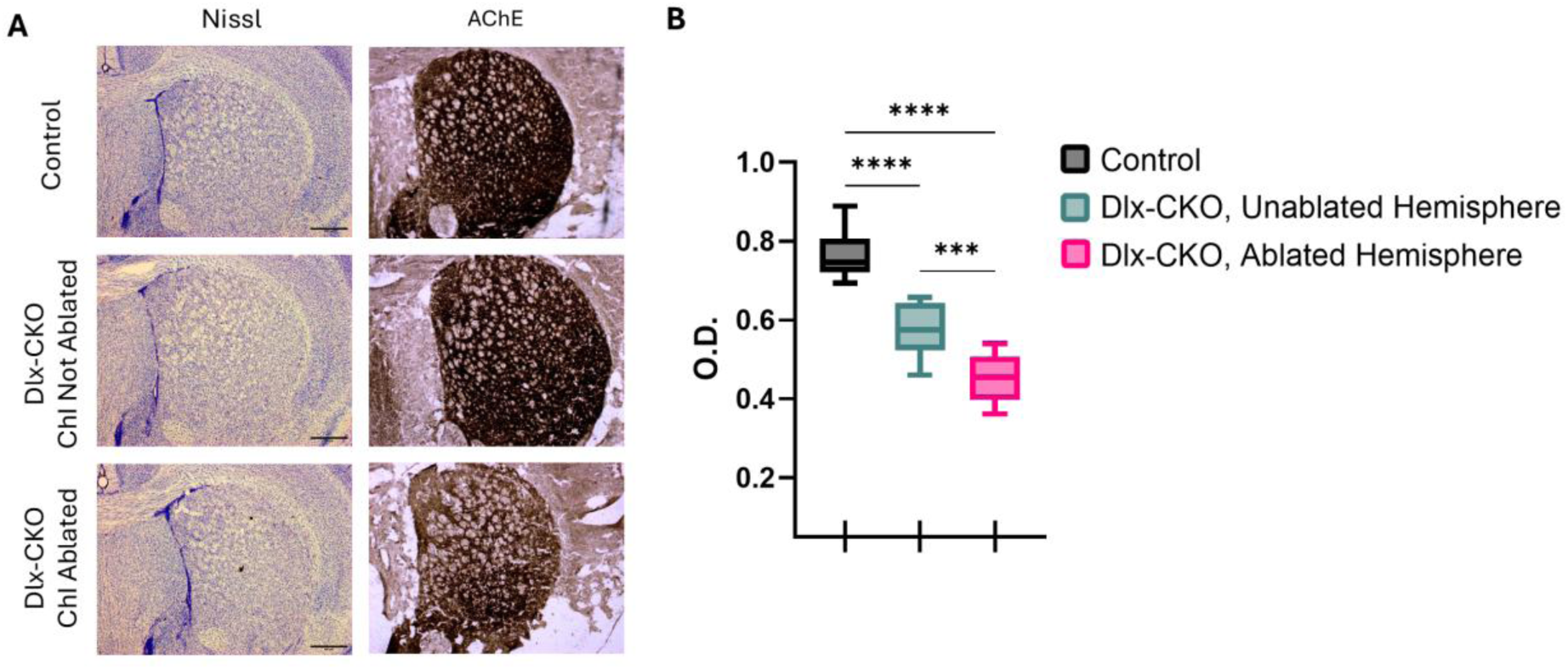
Effects of striatal ChI ablation on acetylcholinesterase levels (A) Representative images of Nissl stain (left panel) and AChE histochemistry (right panel) in the dorsal striatum. Scale bar = 500 μm (B) Optical density measurements of AChE activity staining (One-way ANOVA F_(3, 41)_ = 48.78 p <0.0001; Tukey’s multiple comparisons: Control vs Unablated p <0.0001; Control vs Ablated p <0.0001, Unablated vs Ablated p = 0.0003.

**Figure 6 Supplemental:**
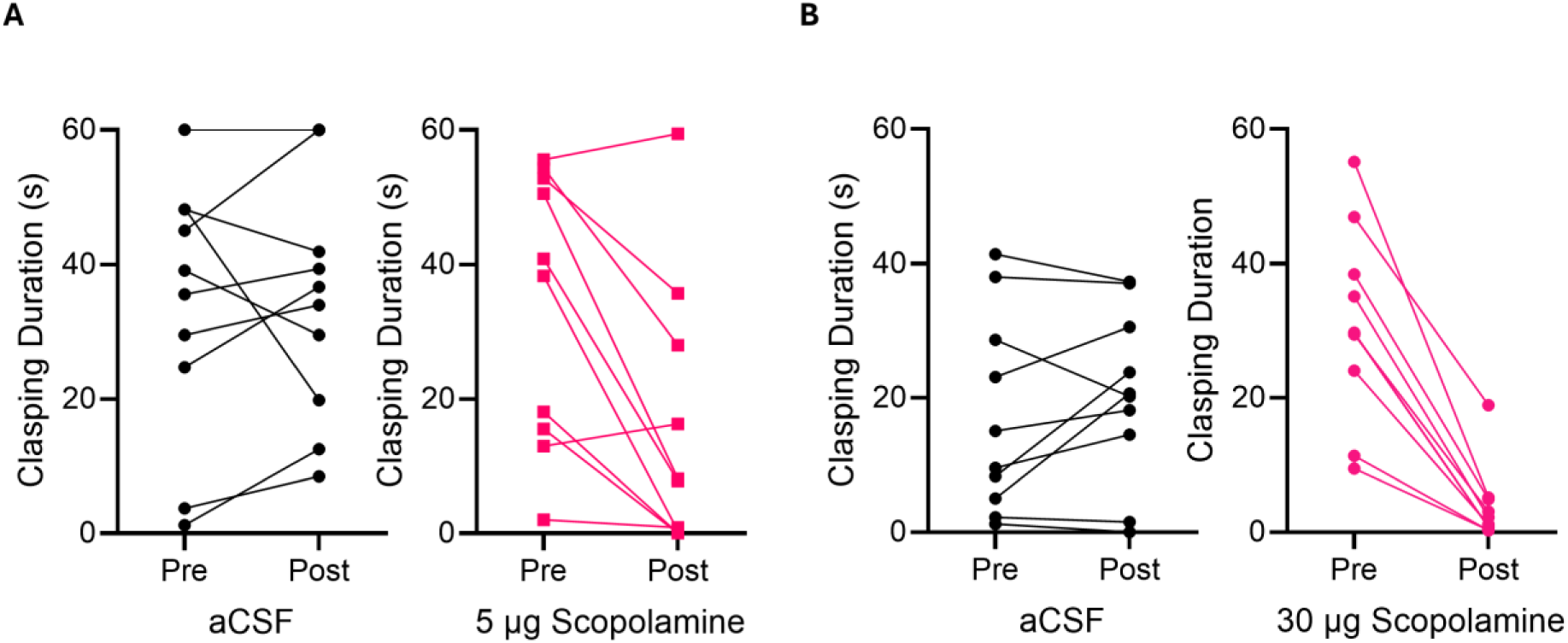
Individual mouse clasping behavior before and after intrastriatal vehicle or scopolamine infusion. (A) Individual mice trials with bilateral vehicle infusion or 5 μg scopolamine. (B) Individual mice trials with bilateral vehicle infusion or 30 μg scopolamine.

**Figure 7 Supplemental:**
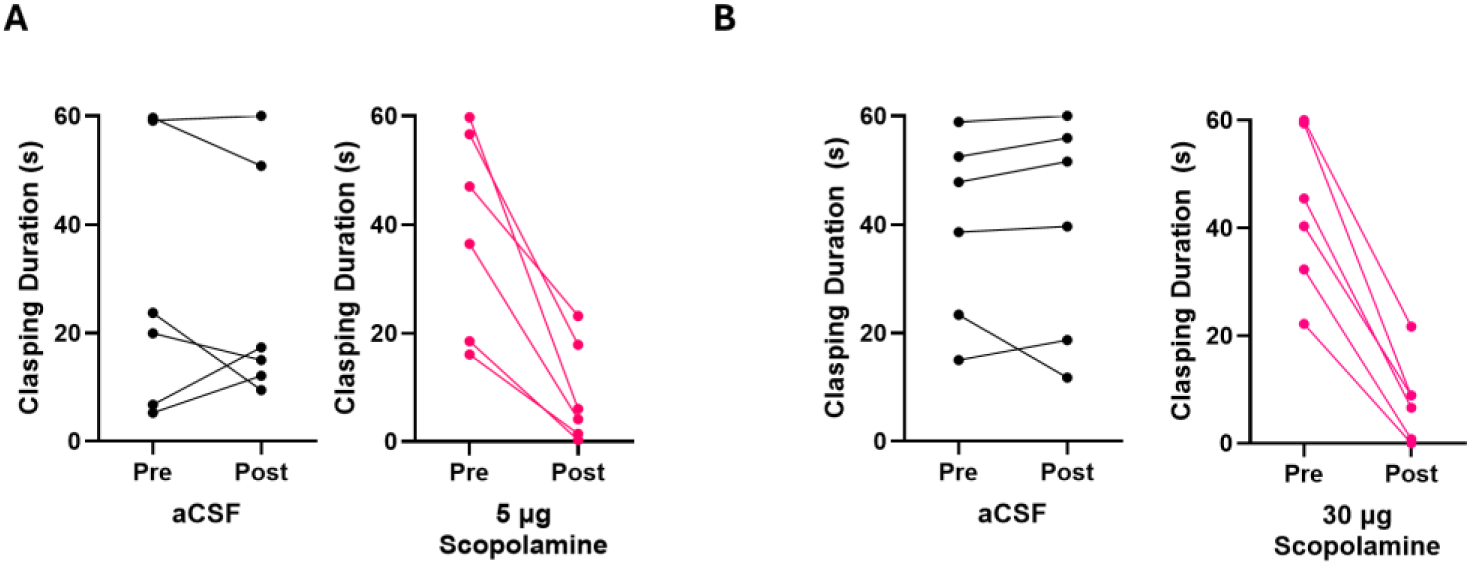
Individual ChI-ablated mouse clasping behavior before and after intrastriatal vehicle or scopolamine infusion. (A) Individual mice trials with bilateral vehicle infusion or 5 μg scopolamine in ChI-ablated Dlx-CKO mice. (B) Individual mice trials with bilateral vehicle infusion or 30 μg scopolamine in ChI-ablated Dlx-CKO mice.

## References

1. Albanese, A., et al., Phenomenology and classification of dystonia: a consensus update. Mov Disord, 2013. 28(7): p. 863–73.

2. Weisheit, C.E., S.S. Pappas, and W.T. Dauer, Inherited dystonias: clinical features and molecular pathways. Handb Clin Neurol, 2018. 147: p. 241–254.

3. Bressman, S.B., et al., The DYT1 phenotype and guidelines for diagnostic testing. Neurology, 2000. 54(9): p. 1746–52.

4. Ozelius, L.J., et al., The early-onset torsion dystonia gene (DYT1) encodes an ATP-binding protein. Nat Genet, 1997. 17(1): p. 40–8.

5. Goodchild, R.E., C.E. Kim, and W.T. Dauer, Loss of the dystonia-associated protein torsinA selectively disrupts the neuronal nuclear envelope. Neuron, 2005. 48(6): p. 923–32.

6. Demircioglu, F.E., et al., Structures of TorsinA and its disease-mutant complexed with an activator reveal the molecular basis for primary dystonia. Elife, 2016. 5.

7. Naismith, T.V., S. Dalal, and P.I. Hanson, Interaction of torsinA with its major binding partners is impaired by the dystonia-associated DeltaGAG deletion. J Biol Chem, 2009. 284(41): p. 27866–27874.

8. Zhao, C., et al., Regulation of Torsin ATPases by LAP1 and LULL1. Proc Natl Acad Sci U S A, 2013. 110(17): p. E1545–54.

9. Zhu, L., et al., A unique redox-sensing sensor II motif in TorsinA plays a critical role in nucleotide and partner binding. J Biol Chem, 2010. 285(48): p. 37271–80.

10. Mazere, J., et al., Striatal and cerebellar vesicular acetylcholine transporter expression is disrupted in human DYT1 dystonia. Brain, 2021. 144(3): p. 909–923.

11. Fan, Y., et al., DYT-TOR1A dystonia: an update on pathogenesis and treatment. Front Neurosci, 2023. 17: p. 1216929.

12. Poppi, L.A., et al., Recurrent Implication of Striatal Cholinergic Interneurons in a Range of Neurodevelopmental, Neurodegenerative, and Neuropsychiatric Disorders. Cells, 2021. 10(4).

13. Burke, R.E., S. Fahn, and C.D. Marsden, Torsion dystonia: a double-blind, prospective trial of high-dosage trihexyphenidyl. Neurology, 1986. 36(2): p. 160–4.

14. Pappas, S.S., et al., Forebrain deletion of the dystonia protein torsinA causes dystonic-like movements and loss of striatal cholinergic neurons. Elife, 2015. 4: p. e08352.

15. Liu, Y., et al., Alteration of the cholinergic system and motor deficits in cholinergic neuron-specific Dyt1 knockout mice. Neurobiol Dis, 2021. 154: p. 105342.

16. Eskow Jaunarajs, K.L., et al., Striatal cholinergic dysfunction as a unifying theme in the pathophysiology of dystonia. Prog Neurobiol, 2015. 127-128: p. 91–107.

17. Wilkes, B.J., et al., Cell-specific effects of Dyt1 knock-out on sensory processing, network-level connectivity, and motor deficits. Exp Neurol, 2021. 343: p. 113783.

18. Pisani, A., et al., Altered responses to dopaminergic D2 receptor activation and N-type calcium currents in striatal cholinergic interneurons in a mouse model of DYT1 dystonia. Neurobiol Dis, 2006. 24(2): p. 318–25.

19. Sciamanna, G., et al., Cholinergic dysfunction alters synaptic integration between thalamostriatal and corticostriatal inputs in DYT1 dystonia. J Neurosci, 2012. 32(35): p. 11991–2004.

20. Yokoi, F., et al., Decreased number of striatal cholinergic interneurons and motor deficits in dopamine receptor 2-expressing-cell-specific Dyt1 conditional knockout mice. Neurobiol Dis, 2020. 134: p. 104638.

21. Ribot, B., et al., Intra-putaminal muscarinic receptor agonist infusion induces a dystonic phenotype in non-human primates. Brain, 2025. 148(12): p. 4532–4547.

22. Gemperli, K., et al., Chronic Striatal Cholinergic Interneuron Excitation Causes Cerebral Palsy-Related Dystonic Behavior in Mice. Ann Neurol, 2025. 98(4): p. 726–740.

23. Li, J., et al., TorsinA restoration in a mouse model identifies a critical therapeutic window for DYT1 dystonia. J Clin Invest, 2021. 131(6).

24. Li, J., et al., CNS critical periods: implications for dystonia and other neurodevelopmental disorders. JCI Insight, 2021. 6(4).

25. Tepper, J.M., et al., Postnatal development of the rat neostriatum: electrophysiological, light- and electron-microscopic studies. Dev Neurosci, 1998. 20(2-3): p. 125–45.

26. McGuirt, A.F., et al., Coordinated Postnatal Maturation of Striatal Cholinergic Interneurons and Dopamine Release Dynamics in Mice. J Neurosci, 2021. 41(16): p. 3597–3609.

27. Plotkin, J.L., et al., Functional and molecular development of striatal fast-spiking GABAergic interneurons and their cortical inputs. Eur J Neurosci, 2005. 22(5): p. 1097–108.

28. Krajeski, R.N., et al., Dynamic postnatal development of the cellular and circuit properties of striatal D1 and D2 spiny projection neurons. J Physiol, 2019. 597(21): p. 5265–5293.

29. Gould, E., N.J. Woolf, and L.L. Butcher, Postnatal development of cholinergic neurons in the rat: I. Forebrain. Brain Res Bull, 1991. 27(6): p. 767–89.

30. Lieberman, O.J., et al., Dopamine Triggers the Maturation of Striatal Spiny Projection Neuron Excitability during a Critical Period. Neuron, 2018. 99(3): p. 540–554 e4.

31. Phelps, P.E., D.R. Brady, and J.E. Vaughn, The generation and differentiation of cholinergic neurons in rat caudate-putamen. Brain Res Dev Brain Res, 1989. 46(1): p. 47–60.

32. Knowles, R., N. Dehorter, and T. Ellender, From Progenitors to Progeny: Shaping Striatal Circuit Development and Function. J Neurosci, 2021. 41(46): p. 9483–9502.

33. Lozovaya, N., S. Eftekhari, and C. Hammond, The early excitatory action of striatal cholinergic-GABAergic microcircuits conditions the subsequent GABA inhibitory shift. Commun Biol, 2023. 6(1): p. 723.

34. Ding, J.B., et al., Thalamic gating of corticostriatal signaling by cholinergic interneurons. Neuron, 2010. 67(2): p. 294–307.

35. Higley, M.J., G.J. Soler-Llavina, and B.L. Sabatini, Cholinergic modulation of multivesicular release regulates striatal synaptic potency and integration. Nat Neurosci, 2009. 12(9): p. 1121–8.

36. Zhang, W., et al., Multiple muscarinic acetylcholine receptor subtypes modulate striatal dopamine release, as studied with M1-M5 muscarinic receptor knock-out mice. J Neurosci, 2002. 22(15): p. 6347–52.

37. Goldberg, J.A., J.B. Ding, and D.J. Surmeier, Muscarinic modulation of striatal function and circuitry. Handb Exp Pharmacol, 2012(208): p. 223–41.

38. Oldenburg, I.A. and J.B. Ding, Cholinergic modulation of synaptic integration and dendritic excitability in the striatum. Curr Opin Neurobiol, 2011. 21(3): p. 425–32.

39. Bonsi, P., et al., Loss of muscarinic autoreceptor function impairs long-term depression but not long-term potentiation in the striatum. J Neurosci, 2008. 28(24): p. 6258–63.

40. Goldberg, J.A. and J.N. Reynolds, Spontaneous firing and evoked pauses in the tonically active cholinergic interneurons of the striatum. Neuroscience, 2011. 198: p. 27–43.

41. McGuirt, A., I. Pigulevskiy, and D. Sulzer, Developmental regulation of thalamus-driven pauses in striatal cholinergic interneurons. iScience, 2022. 25(11): p. 105332.

42. Pisani, A., et al., Re-emergence of striatal cholinergic interneurons in movement disorders. Trends Neurosci, 2007. 30(10): p. 545–53.

43. Ding, Y., et al., Enhanced striatal cholinergic neuronal activity mediates L-DOPA-induced dyskinesia in parkinsonian mice. Proc Natl Acad Sci U S A, 2011. 108(2): p. 840–5.

44. Surmeier, D.J. and A.M. Graybiel, A feud that wasn’t: acetylcholine evokes dopamine release in the striatum. Neuron, 2012. 75(1): p. 1–3.

45. Lester, D.B., T.D. Rogers, and C.D. Blaha, Acetylcholine-dopamine interactions in the pathophysiology and treatment of CNS disorders. CNS Neurosci Ther, 2010. 16(3): p. 137–62.

46. Yellajoshyula, D., et al., Genetic evidence of aberrant striatal synaptic maturation and secretory pathway alteration in a dystonia mouse model. Dystonia, 2022. 1.

47. Pappas, S.S., et al., A cell autonomous torsinA requirement for cholinergic neuron survival and motor control. Elife, 2018. 7.

48. Bertran-Gonzalez, J., et al., Striatal cholinergic interneurons display activity-related phosphorylation of ribosomal protein S6. PLoS One, 2012. 7(12): p. e53195.

49. Fahn, S., High dosage anticholinergic therapy in dystonia. Neurology, 1983. 33(10): p. 1255–61.

50. Greene, P., H. Shale, and S. Fahn, Analysis of open-label trials in torsion dystonia using high dosages of anticholinergics and other drugs. Mov Disord, 1988. 3(1): p. 46–60.

51. Phelps, P.E., C.R. Houser, and J.E. Vaughn, Immunocytochemical localization of choline acetyltransferase within the rat neostriatum: a correlated light and electron microscopic study of cholinergic neurons and synapses. J Comp Neurol, 1985. 238(3): p. 286–307.

52. Woolf, N.J. and L.L. Butcher, Cholinergic neurons in the caudate-putamen complex proper are intrinsically organized: a combined Evans blue and acetylcholinesterase analysis. Brain Res Bull, 1981. 7(5): p. 487–507.

53. Dautan, D., et al., A major external source of cholinergic innervation of the striatum and nucleus accumbens originates in the brainstem. J Neurosci, 2014. 34(13): p. 4509–18.

54. Dautan, D., et al., Extrinsic Sources of Cholinergic Innervation of the Striatal Complex: A Whole-Brain Mapping Analysis. Front Neuroanat, 2016. 10: p. 1.

55. Satoh, K., et al., Ultrastructural observations of the cholinergic neuron in the rat striatum as identified by acetylcholinesterase pharmacohistochemistry. Neuroscience, 1983. 10(4): p. 1121–36.

56. Holler, T., et al., Differences in the developmental expression of the vesicular acetylcholine transporter and choline acetyltransferase in the rat brain. Neurosci Lett, 1996. 212(2): p. 107–10.

57. Graybiel, A.M. and C.W. Ragsdale, Jr., Histochemically distinct compartments in the striatum of human, monkeys, and cat demonstrated by acetylthiocholinesterase staining. Proc Natl Acad Sci U S A, 1978. 75(11): p. 5723–6.

58. Dorje, F., et al., Antagonist binding profiles of five cloned human muscarinic receptor subtypes. J Pharmacol Exp Ther, 1991. 256(2): p. 727–33.

59. Levey, A.I., et al., Identification and localization of muscarinic acetylcholine receptor proteins in brain with subtype-specific antibodies. J Neurosci, 1991. 11(10): p. 3218–26.

60. Lebois, E.P., et al., Muscarinic receptor subtype distribution in the central nervous system and relevance to aging and Alzheimer’s disease. Neuropharmacology, 2018. 136(Pt C): p. 362–373.

61. Yan, Z., J. Flores-Hernandez, and D.J. Surmeier, Coordinated expression of muscarinic receptor messenger RNAs in striatal medium spiny neurons. Neuroscience, 2001. 103(4): p. 1017–24.

62. Abudukeyoumu, N., et al., Cholinergic modulation of striatal microcircuits. Eur J Neurosci, 2019. 49(5): p. 604–622.

63. Weiner, D.M., A.I. Levey, and M.R. Brann, Expression of muscarinic acetylcholine and dopamine receptor mRNAs in rat basal ganglia. Proc Natl Acad Sci U S A, 1990. 87(18): p. 7050–4.

64. Vilaro, M.T., J.M. Palacios, and G. Mengod, Localization of m5 muscarinic receptor mRNA in rat brain examined by in situ hybridization histochemistry. Neurosci Lett, 1990. 114(2): p. 154–9.

65. Hersch, S.M., et al., Distribution of m1-m4 muscarinic receptor proteins in the rat striatum: light and electron microscopic immunocytochemistry using subtype-specific antibodies. J Neurosci, 1994. 14(5 Pt 2): p. 3351–63.

66. Calabresi, P., et al., Blockade of M2-like muscarinic receptors enhances long-term potentiation at corticostriatal synapses. Eur J Neurosci, 1998. 10(9): p. 3020–3.

67. Bernard, V., E. Normand, and B. Bloch, Phenotypical characterization of the rat striatal neurons expressing muscarinic receptor genes. J Neurosci, 1992. 12(9): p. 3591–600.

68. Zhang, W., et al., Characterization of central inhibitory muscarinic autoreceptors by the use of muscarinic acetylcholine receptor knock-out mice. J Neurosci, 2002. 22(5): p. 1709–17.

69. Yan, Z. and D.J. Surmeier, Muscarinic (m2/m4) receptors reduce N- and P-type Ca2+ currents in rat neostriatal cholinergic interneurons through a fast, membrane-delimited, G-protein pathway. J Neurosci, 1996. 16(8): p. 2592–604.

70. Calabresi, P., et al., Muscarinic IPSPs in rat striatal cholinergic interneurones. J Physiol, 1998. 510 (Pt 2)(Pt 2): p. 421–7.

71. Chen, L., M. Chatterjee, and J.Y. Li, The mouse homeobox gene Gbx2 is required for the development of cholinergic interneurons in the striatum. J Neurosci, 2010. 30(44): p. 14824–34.

72. Semba, K., S.R. Vincent, and H.C. Fibiger, Different times of origin of choline acetyltransferase- and somatostatin-immunoreactive neurons in the rat striatum. J Neurosci, 1988. 8(10): p. 3937–44.

73. Casarosa, P., et al., The constitutive activity of the human muscarinic M3 receptor unmasks differences in the pharmacology of anticholinergics. J Pharmacol Exp Ther, 2010. 333(1): p. 201–9.

74. Spalding, T.A. and E.S. Burstein, Constitutive activity of muscarinic acetylcholine receptors. J Recept Signal Transduct Res, 2006. 26(1-2): p. 61–85.

75. Bernard, V., et al., Subcellular redistribution of m2 muscarinic acetylcholine receptors in striatal interneurons in vivo after acute cholinergic stimulation. J Neurosci, 1998. 18(23): p. 10207–18.

76. Bernard, V., A.I. Levey, and B. Bloch, Regulation of the subcellular distribution of m4 muscarinic acetylcholine receptors in striatal neurons in vivo by the cholinergic environment: evidence for regulation of cell surface receptors by endogenous and exogenous stimulation. J Neurosci, 1999. 19(23): p. 10237–49.

77. Lim, S.A., U.J. Kang, and D.S. McGehee, Striatal cholinergic interneuron regulation and circuit effects. Front Synaptic Neurosci, 2014. 6: p. 22.

78. Gonzales, K.K. and Y. Smith, Cholinergic interneurons in the dorsal and ventral striatum: anatomical and functional considerations in normal and diseased conditions. Ann N Y Acad Sci, 2015. 1349(1): p. 1–45.

79. Koos, T. and J.M. Tepper, Dual cholinergic control of fast-spiking interneurons in the neostriatum. J Neurosci, 2002. 22(2): p. 529–35.

80. Calabresi, P., et al., Activation of M1-like muscarinic receptors is required for the induction of corticostriatal LTP. Neuropharmacology, 1999. 38(2): p. 323–6.

81. Wang, Z., et al., Dopaminergic control of corticostriatal long-term synaptic depression in medium spiny neurons is mediated by cholinergic interneurons. Neuron, 2006. 50(3): p. 443–52.

82. Bolden, C., B. Cusack, and E. Richelson, Antagonism by antimuscarinic and neuroleptic compounds at the five cloned human muscarinic cholinergic receptors expressed in Chinese hamster ovary cells. J Pharmacol Exp Ther, 1992. 260(2): p. 576–80.

83. Moehle, M.S., et al., Discovery of the First Selective M(4) Muscarinic Acetylcholine Receptor Antagonists with in Vivo Antiparkinsonian and Antidystonic Efficacy. ACS Pharmacol Transl Sci, 2021. 4(4): p. 1306–1321.

84. Sheffler, D.J., et al., A novel selective muscarinic acetylcholine receptor subtype 1 antagonist reduces seizures without impairing hippocampus-dependent learning. Mol Pharmacol, 2009. 76(2): p. 356–68.

85. Shirey, J.K., et al., A selective allosteric potentiator of the M1 muscarinic acetylcholine receptor increases activity of medial prefrontal cortical neurons and restores impairments in reversal learning. J Neurosci, 2009. 29(45): p. 14271–86.

86. Maltese, M., et al., Anticholinergic drugs rescue synaptic plasticity in DYT1 dystonia: role of M1 muscarinic receptors. Mov Disord, 2014. 29(13): p. 1655–65.

87. Downs, A.M., et al., Blockade of M4 muscarinic receptors on striatal cholinergic interneurons normalizes striatal dopamine release in a mouse model of TOR1A dystonia. Neurobiol Dis, 2022. 168: p. 105699.

88. Lange, L.M., et al., Genotype-Phenotype Relations for Isolated Dystonia Genes: MDSGene Systematic Review. Mov Disord, 2021. 36(5): p. 1086–1103.

89. Balint, B., et al., Dystonia. Nat Rev Dis Primers, 2018. 4(1): p. 25.

90. Monory, K., et al., The endocannabinoid system controls key epileptogenic circuits in the hippocampus. Neuron, 2006. 51(4): p. 455–66.

91. Franklin, K.B.J. and G. Paxinos, The mouse brain in stereotaxic coordinates. 1997, San Diego: Academic Press. xxii p., 186 p. of plates.

